# TDP1 drives MRE11-dependent and independent end resection during replication stress by TOP2 poisons

**DOI:** 10.64898/2026.05.29.728668

**Authors:** Néstor Aznar, María Camila Gosso, Irene Larripa, Marcela González-Cid, Marcelo de Campos Nebel

## Abstract

DNA-Topoisomerase crosslinks are lesions formed by proteins covalently bound to DNA, and these complexes directly impede transcription and replication, thereby threatening genomic integrity. Tyrosyl-DNA phosphodiesterase 1 (TDP1) and MRE11 remove DNA-protein adducts, but their functional relationship remains unclear. We show TDP1 and MRE11 act epistatically in the removal of etoposide-stimulated topoisomerase IIα cleavage complexes (TOP2αcc) in S/G2 phases of the cell cycle. Etoposide-stimulated TOP2αcc formed ahead of and behind replication forks, inhibiting fork progression. Consistently, replication fork progression was decreased in cells lacking TDP1 or upon inhibition of MRE11 nuclease activity, and a similar defect was observed when combining both conditions. Furthermore, TDP1 promotes DNA end resection downstream of MRE11 and facilitates nascent strand degradation at stalled forks independently of MRE11 nuclease activity. Accordingly, combined TDP1 loss and MRE11 inhibition did not exacerbate etoposide hypersensitivity. We provide evidence that TDP1 stimulates replication-coupled removal of TOP2αcc by facilitating DNA end processing through both MRE11-dependent and –independent pathways. Overall, our results suggest TDP1 promotes nucleolytic excision of TOP2α-mediated replication blocks, in addition to its canonical hydrolase activity.

## INTRODUCTION

The double-stranded DNA molecules contain the genetic information of living cells. DNA metabolic transactions such as transcription or replication require the unwinding of the double-stranded DNA molecules to take place. The outcome of these nuclear activities is the alteration of the topological state of the DNA. Topoisomerases are essential enzymes involved in the regulation of DNA topology playing critical roles in various cellular processes through the introduction of transient breaks in DNA (Nitiss, 2009a). Mammalian type IIA topoisomerases (TOP2α and TOP2β) utilize transient double-strand breaks (DSB) to mediate the passage of one DNA segment through another. This enzymatic activity is critical for regulating DNA supercoiling and resolving catenanes in DNA molecules. TOP2α is primarily associated with DNA replication and chromosome segregation in dividing cells. TOP2β has a particular role in transcription and chromatin organization (Nitiss, 2009a, Pommier, Leo et al., 2010). In any case, TOP2 utilizes ATP hydrolysis to cleave, pass, and religate DNA strands (Wang, 1996). The cleavage step of TOP2 generates a transient covalent bond between a tyrosine residue from its active site and the 5’end of the DNA backbone, also called topoisomerase II cleavage complex (TOP2cc) (Nitiss, 2009a, Wang, 2002).

TOP2 is the target of a broad range of anticancer drugs, including several epipodophyllotoxins, anthracyclines, and anthracenediones (Nitiss, 2009b, Pommier et al., 2010). The paradigm of TOP2 poison is etoposide (ETO), utilized in the treatment of both solid tumors and onco-hematological malignancies (Bailly, 2023, Vann, Oviatt et al., 2021, Zhang, Gou et al., 2021). TOP2 poisons stabilize TOP2cc, increasing its half-life and leading to protein-linked DNA double-strand break formation. TOP2-mediated DNA damage, an enzymatic type of DNA-protein crosslink (DPC), requires both the removal of the bulky protein as well as the repair of the resulting DNA DSB (Sun, Miller Jenkins et al., 2020). The repair of TOP2-DPCs requires the coordinated action of proteases, nucleases and hydrolases to prevent genome instability (Weickert & Stingele, 2022).

Tyrosyl-DNA phosphodiesterase 1 (TDP1) and 2 (TDP2) are specialized hydrolases involved in the cleavage of phosphodiester bonds between the DNA and proteins. Pathogenic mutations in TDP1 are associated to the neurological condition of spinocerebellar ataxia with axonal neuropathy (SCAN1) (Takashima, Boerkoel et al., 2002), while mutations in TDP2 cause the spinocerebellar ataxia autosomal recessive 23 (SCAR23) (Zagnoli-Vieira, Bruni et al., 2018). TDP1 has a broad variety of substrates and potent phosphodiesterase activity carried out by its two catalytic HKD motifs located in close proximity in the C-terminus portion of the protein (Interthal, Pouliot et al., 2001). In straight contrast, TDP2 shows a specific phosphodiesterase activity but limited to phosphotyrosyl bonds through four conserved motifs (TWN, LQE, GDXN and SDH) which are critical for its activity (Cortes Ledesma, El Khamisy et al., 2009). TDP2 primary acts on proteins bound to DNA with accessible 5’ ends, while TDP1 targets proteins tyrosyl-linked to 3’ DNA termini (Pommier, Huang et al., 2014). Interestingly, both enzymes show overlapping activities on the removal of topoisomerase adducts (Borda, Palmitelli et al., 2015, Murai, Huang et al., 2012, Tsuda, Kitamasu et al., 2020).

Meiotic Recombination 11 (MRE11), a nuclease that interacts with RAD50 and NBS1, forms the MRN complex which plays a central role in sensing DSB and initiates the DNA end-resection process, which in turn triggers the repair by homologous recombination (HR) (Buis, Wu et al., 2008, Paull & Gellert, 1998). Mutations in the *MRE11A* gene are responsible for an ataxia telangiectasia-like disorder 1 (ATLD1) (Sedghi, Salari et al., 2018). MRE11 harbors four N-terminal phosphodiesterase motifs that constitute its catalytic domain (Hopfner, Karcher et al., 2001). Both the endo– and exonuclease activities of MRE11 are required for DNA end resection (Garcia, Phelps et al., 2011, Shibata, Moiani et al., 2014, Wang, Daley et al., 2017). MRE11’s nuclease activity enables HR by generating 3’ single-stranded tails at DSBs and by facilitating the recruitment of long-range nucleases EXO1 and DNA2 (Mimitou & Symington, 2008, Zhu, Chung et al., 2008).

In lower eukaryotes, protein adducts at DNA ends stimulate the MRN complex’s endonuclease activity, resulting in the cleavage of these adducts (Cannavo & Cejka, 2014, Deshpande, Lee et al., 2016). Similarly, in human cells, MRE11 plays a crucial role in removing TOP2 from DNA, a process that prevents the accumulation of stalled TOP2 complexes caused by abortive catalysis or exposure to TOP2 poisons (Hoa, Shimizu et al., 2016b, Lee, Padget et al., 2012). The strategy of using its endo– and exonuclease activities to remove protein adducts from DNA ends is conserved throughout the evolution, extending from phages to both pro– and eukaryotes (Connelly, de Leau et al., 2003, Lee et al., 2012, Neale, Pan et al., 2005, Stohr & Kreuzer, 2001).

The multiple DPC repair pathways reflect the significant challenge they pose to genomic integrity. However, the coordination among these repair factors and the pathway selection are still poorly understood.

Here, we show that the removal of DNA-bound TOP2α by TDP1 and MRE11 is determined by an epistatic relationship between these enzymes during the S/G2 phases of the cell cycle. Furthermore, we find that TOP2α-mediated cleavage complexes form on both sides of the replication fork and identify an additional role for TDP1 in promoting DNA end-resection following replicative stress.

## RESULTS

### TDP1 deficient cells accumulate increased amounts of ETO-induced TOP2αcc during S and G2 phase of the cell cycle

We have previously demonstrated that lack of TDP1 results in an increased accumulation of ETO-induced TOP2αcc (Borda et al., 2015). Thus, we investigated whether the increased levels of TOP2αcc by lack of TDP1 differs across cell cycle phases to identify potential mechanisms affected. As shown in Fig. 1A-B, HeLa TDP1 deficient cells (TDP1kd) treated with ETO 17 μM accumulated significantly higher levels of TOP2αcc in S/G2 phases of the cell cycle with respect to non-silencing control cells (NS) (P<0.03). As expected, the increase in TOP2αcc levels observed in TDP1kd cells was milder than those found in TDP2 deficient cells (TDP2kd) (Fig. 1C). Next, we analyzed the formation of DSB through γH2AX detection at the same time in the different cell lines. As shown in Fig. 1D-E, ETO-treatment increased the γH2AX intensity in both NS and TDP1kd cells throughout the cell cycle (*P*<0.0009 and *P*<0.0004, respectively). However, γH2AX levels in S/G2 phases showed no significant differences between NS and TDP1kd cells. Interestingly, TDP2kd cells (Fig. 1F) showed a significant increase in ETO-induced γH2AX levels just in G1 cells (*P*<0.02).

**Figure 1:**
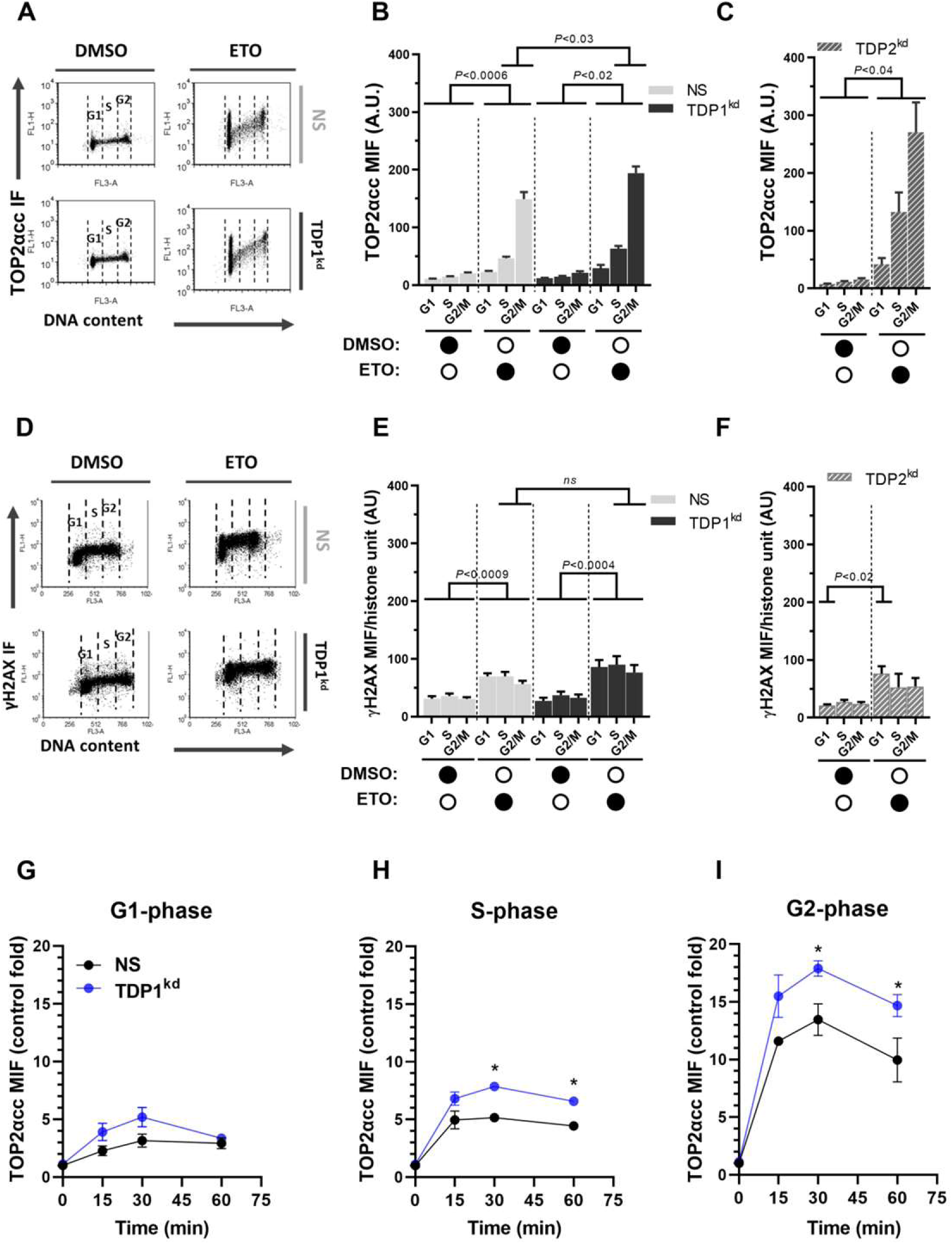
TDP1 deficient cells accumulate ETO-induced TOP2αcc in S/G2 phase of the cell cycle. (A) Representative dot plot images of TOP2αcc of NS and TDP1kd cells after exposure to ETO 17 µM by 1h. Dotted-lines separate the different phases of the cell cycle. (B) Bar graph showing TOP2αcc median intensity of fluorescence (MIF) induced by ETO in the different cell cycle phases of NS and TDP1kd cells. (C) Bar graph showing ETO-induced TOP2αcc MIF in TDP2kd cells. (D) Dot-plot images of γH2AX induced by ETO 17µM after exposure by 1h in NS and TDP1kd cells. Dotted-lines separate the different cell cycle phases. (E) Bar graph showing γH2AX median intensity of fluorescence (MIF) per histone unit induced by 1h exposure to ETO 17 µM in NS and TDP1kd cells. *ns*= no significant differences. (F) Bar graph showing ETO-induced γH2AX MIF per histone unit in TDP2kd cells. (G-I) TOP2αcc accumulation of NS and TDP1kd cells after treatment with ETO 17µM by different exposure times in G1 phase (G), S phase (H), and G2 phase cells (I). \**P*<0.05. (B, C, E, F, G-I) Error bars indicate s.e.m.

In order to determine whether the increased detection of ETO-induced TOP2αcc in TDP1kd cells was due to an accumulation of these complexes, we examined their formation over the time. As shown in Fig. 1G-I, TOP2αcc accumulated in a time-dependent manner and was significantly higher in S/G2 phases of TDP1kd cells compared to NS cells (*P*<0.05).

Thus, our results show that lack of TDP1 cause an accumulation of ETO-induced TOP2αcc in the S and G2 phases of the cell cycle. However, the increased levels of TOP2αcc were not accompanied by a diminished amount of γH2AX signals at the time point and dose analyzed.

### Transcription inhibition does not contribute to increase ETO-induced TOP2αcc levels or to reduce γH2AX signals

Previous studies suggested that drug-stabilized TOP2cc requires at least partially a transcription-dependent processing (Fan, Peng et al., 2008, Tammaro, Barr et al., 2013, Zhang, Lyu et al., 2006). Moreover, it has been proposed that collision of elongating RNA polymerase II with TOP2cc results in TOP2 degradation and DSB signaling (Yan, Tammaro et al., 2016).

To investigate if transcription bubbles are involved in the collision with TOP2α-mediated cleavage complexes following a short-term treatment with ETO, we analyzed TOP2αcc and γH2AX levels using the transcription inhibitor 5,6-dichloro-1-beta-D-ribofuranosylbenzimidazole (DRB). Cell cultures were pre-treated with DRB before a treatment with a high dose of ETO (250 μM). As shown in Fig. 2A-B, transcriptional inhibition by DRB promoted a non-significant increase of ETO-induced TOP2αcc levels throughout the cell cycle, in both NS and TDP1kd cells.

**Figure 2:**
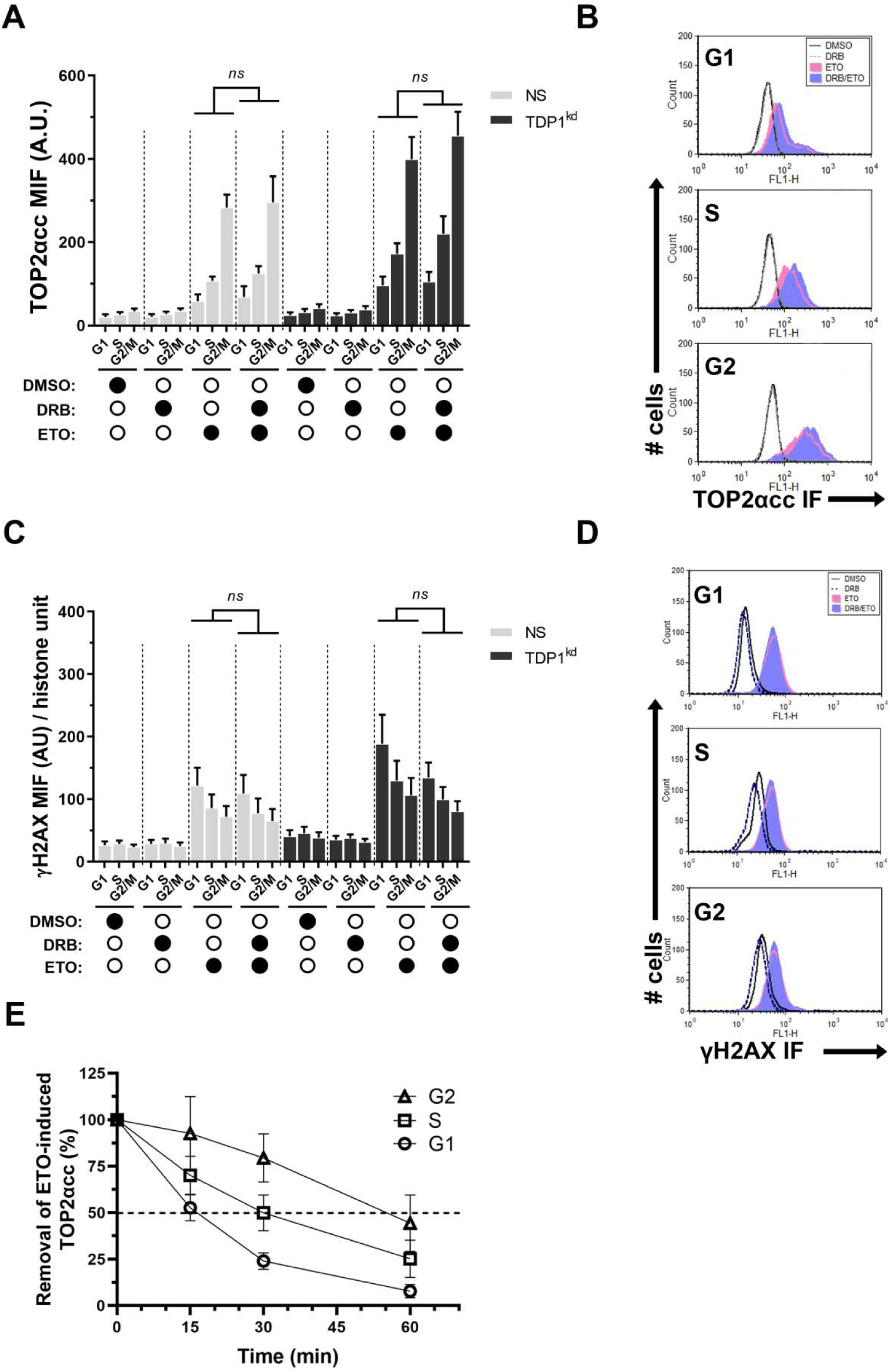
The accumulation of ETO-induced TOP2αcc and the levels of γH2AX are both unaltered by transcription inhibition. NS and TDP1kd cells were incubated or not with DRB 150 µM and then treated with ETO 250µM by 1h before the analysis of (A) TOP2αcc median intensity of fluorescence (MIF) or (C) γH2AX MIF in the different cell cycle phases. A.U.= arbitrary units. *ns*= no significant differences. (B) Histogram plots showing levels of TOP2αcc in NS cells in the different cell cycle phases from a representative experiment. (D) Histogram plots showing γH2AX levels from NS cells throughout the cell cycle from a representative experiment. (E) Removal time of ETO-induced TOP2αcc in the different cell cycle phases from NS cells. (A, C, E) Error bars represent s.e.m.

To assess the impact of transcription inhibition on the processing of TOP2cc into DSBs or the collision of RNA polymerase II with them, γH2AX levels were analyzed. As shown in Fig. 2C-D, DRB treatment did not significantly modify ETO-induced γH2AX levels across the cell cycle or cell lines. Similar results were obtained with the unspecific transcriptional inhibitor Actinomycin D (data not shown). Intriguingly, both NS and TDP1kd cells exhibited elevated γH2AX levels in the G1 phase, coinciding with the cell cycle phase in which lower TOP2α activity and TOP2αcc levels are found. Thus, we analyzed in NS cells the time taken to remove TOP2αcc in the different phases of the cell cycle in order to determine whether they are processed at different velocities. As shown in Figure 2E, ETO-stabilized TOP2αcc were removed with different kinetics across cell cycle. The half-life of TOP2αcc was shorter in G1 (∼15 min) compared to S (∼30 min) and G2 (∼55 min) phases.

Overall, our findings suggest that transcription inhibition does not increase ETO-induced TOP2α-mediated cleavage complexes after 1h exposure; thus, the collisions of the transcription machinery with TOP2αcc or a transcription-dependent processing of TOP2αcc does not contribute to DNA DSB formation.

### ETO stabilize TOP2αcc on both sides of the ongoing replication fork

In order to determine whether TOP2αcc stabilized by ETO are formed in the chromatin ahead of the replication fork, we performed pre-treatments with APH before the treatment with ETO 17 μM. APH, a known inhibitor of DNA polymerases α and δ, uncouples the replication forks (Walter & Newport, 2000). This uncoupling allows the replicative helicase to continue unwinding DNA, generating positive supercoiling ahead of the replication fork on the parental strands. Therefore, NS cells were pre-treated with APH for 1 h or left untreated before exposure to ETO for different time points (Fig. 3A). The accumulation of TOP2αcc in the S-phase was subsequently analyzed. Fig. 3B-C shows that while ETO-induced TOP2αcc remained stable from 30 to 60 min (∼5.9 fold of control), the uncoupling of the replication forks by APH increased the complexes induced by ETO from 6.5 to 8.2 fold during the same period of time (*P*=0.0067). This suggests that TOP2αcc stabilized by ETO are being accumulated ahead of the replication forks. A schematic representation is shown in Fig. 3D.

**Figure 3:**
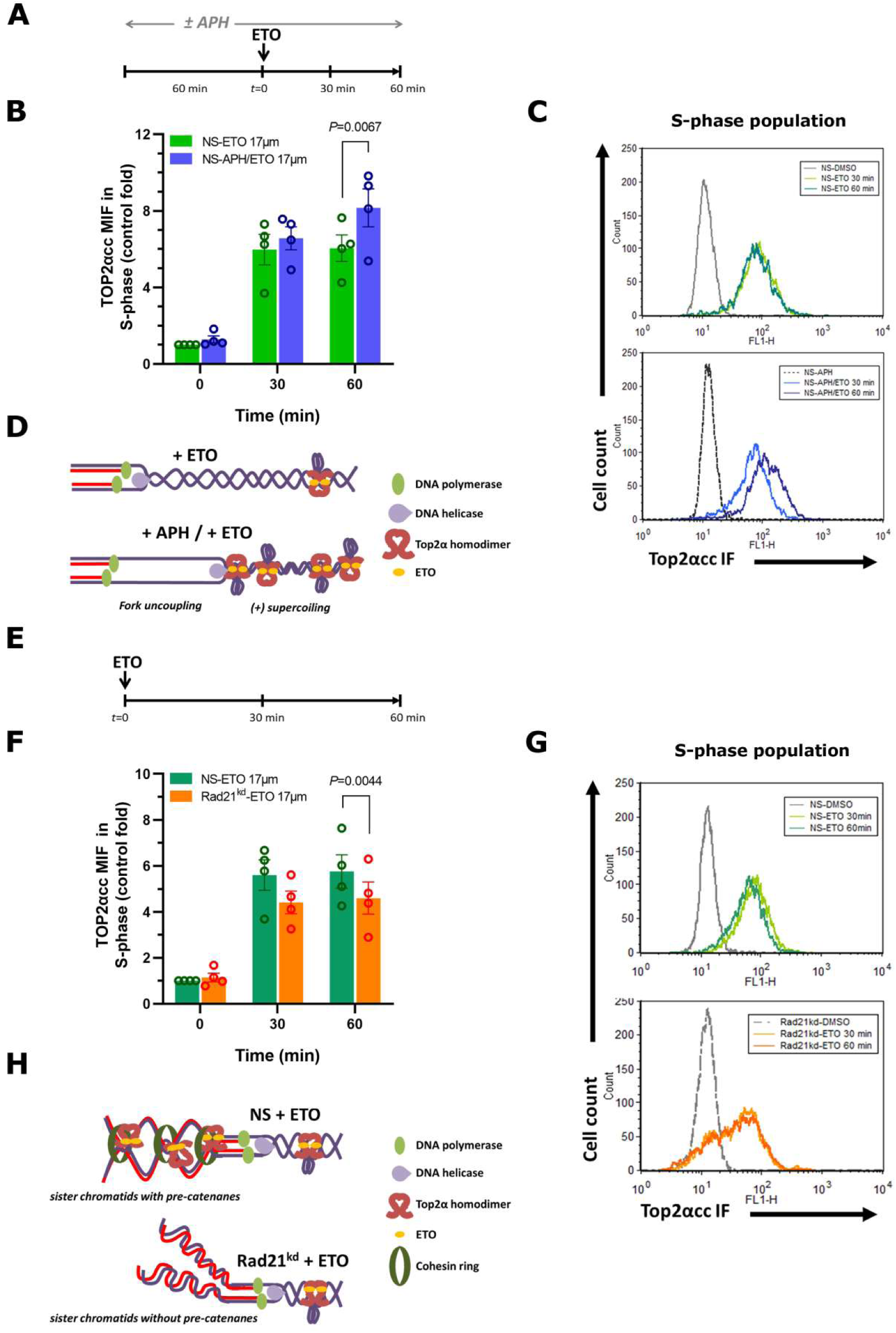
ETO-induced TOP2αcc occur at both sides of the ongoing replication fork. NS cells were pre-treated or not with APH 4 µM by 1h to induce fork uncoupling before adding ETO 17µM by 30 or 60 min. (A) Schematic representation of treatments over the time. (B) Bar graph showing the median intensity of fluorescence (MIF) of ETO-induced TOP2αcc in the S-phase of NS cells. Error bars represent s.e.m. (C) Representative histogram plots of NS cells from an independent experiment. (D) TOP2αcc accumulation ahead of replication forks following fork uncoupling and the associated generation of (+) supercoiling. (E) Schematic representation of treatments over the time. (F) Bar graph showing the median intensity of fluorescence (MIF) of ETO-induced TOP2αcc in the S-phase of NS and Rad21kd cells. Error bars represent s.e.m. (G) Representative histogram plots of NS and Rad21kd cells from an independent experiment. (H) TOP2αcc formation is stimulated behind of the replication fork by the presence of pre-catenanes.

The cohesin complex is a ring-like structure that is essential for sister chromatid cohesion and DNA repair. Within this ring, the RAD21 protein remains bridging other two subunits of the complex, the SMC1 and SMC3 proteins (Dorsett & Strom, 2012). In order to determine whether TOP2α-mediated cleavage complexes are formed behind the ongoing replication fork, we analyzed the accumulation of ETO-induced TOP2αcc in the NS and RAD21kd cell lines. As depicted in Fig. 3E, NS and RAD21kd cells were incubated with ETO 17 μM during different time points. The results are shown in Fig. 3F-G. The accumulation of TOP2αcc induced by ETO in NS cells was similar at 30 and 60 min of treatment (∼5.8 fold of control), as reported above. However, the accumulation of TOP2αcc after 60 min of ETO in RAD21kd cells was significantly lower (∼4.5 fold) than those found in NS control cells (*P*=0.0044). A schematic representation is depicted in Fig. 3H.

Together, our results suggest that in a chromatin context, TOP2αcc induced by ETO accumulate ahead and behind of the ongoing replication forks.

### TDP1 and MRE11 act epistatically to remove ETO-induced TOP2αcc and facilitate replication fork progression

TDP1 and MRE11 participate in the removal of drug-stabilized TOP2αcc. Both nucleases display nucleolytic activity during the S and G2 phases of the cell cycle. To investigate a potential cooperation between MRE11 and TDP1, we first examined the accumulation of ETO-induced TOP2αcc in NS and TDP1kd cells, with or without mre11i pre-treatment. As shown in Fig. 4A, mre11i significantly increased ETO-induced TOP2αcc levels in NS cells during S and G2 phases compared to ETO alone (P<0.05). While ETO alone increased TOP2αcc levels in TDP1kd cells compared to NS cells (P<0.05), no further increase was observed in S and G2 phases of TDP1kd cell line when pre-treated with mre11i and exposed to ETO.

**Figure 4:**
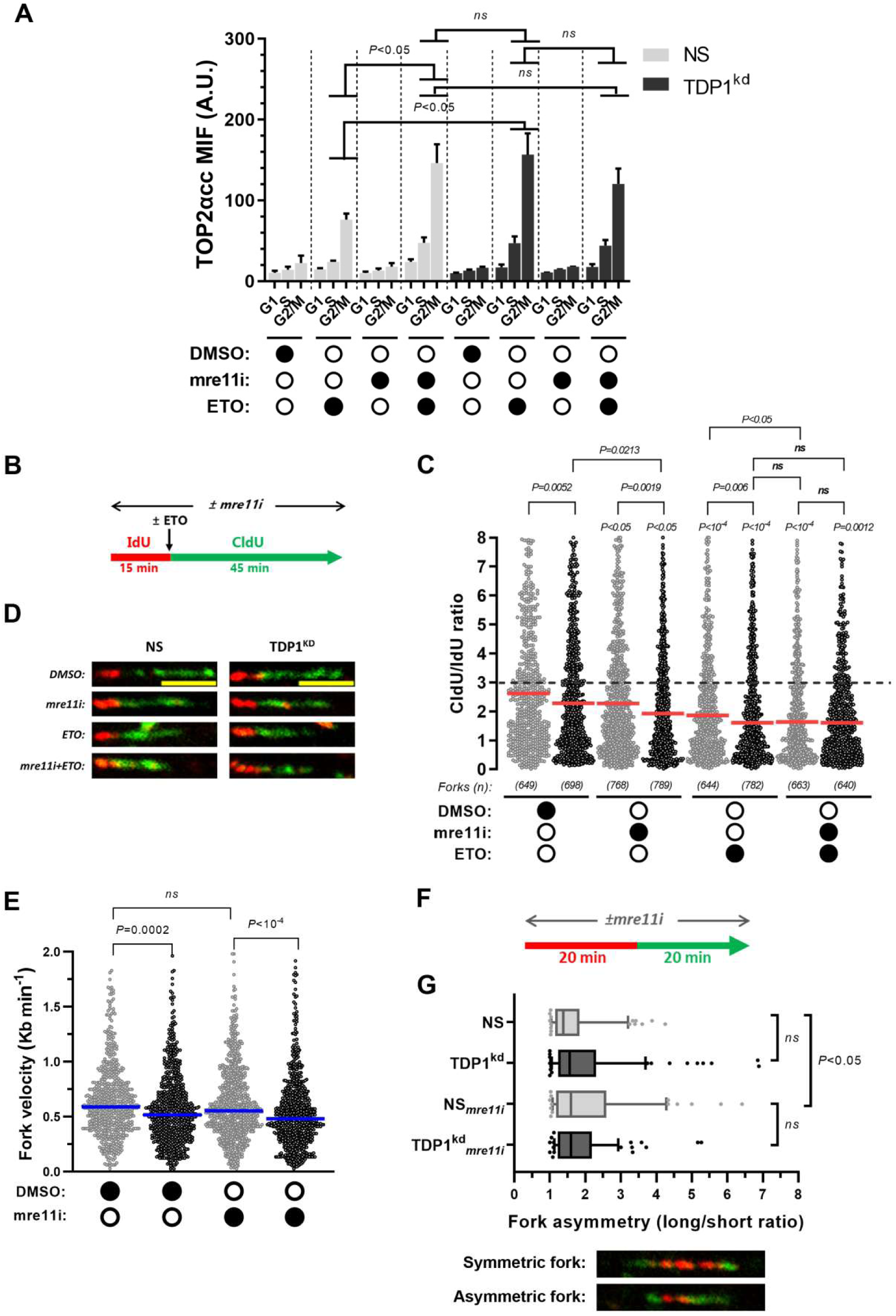
Functional interaction between TDP1 and MRE11 for the removal of TOP2αcc during replication. (A) NS and TDP1kd cells were pretreated or not with mre11i 100 µM by 1h before treatment with ETO 17 µM by an additional hour and analyzed the median intensity of fluorescence (MIF) of ETO-induced TOP2αcc. A.U. = arbitrary units. *ns*= no significant differences. (B) Schematic representation of incubation times with the chemical inhibitors and nucleoside analogues for the evaluation of fork progression. (C) Scatter plot showing fork progression assays performed in NS and TDP1kd cells pretreated or not with mre11i and treated with ETO. *ns*= no significant differences. The red line represents median values of CldU/IdU ratios. (D) Representative fibers of NS and TDP1kd cells with the different treatments. Scale bar= 5 µM. (E-G) Replication fork speed and fork asymmetry. (E) Scatter plot showing the replication fork speed of NS and TDP1kd after incubation with mre11i 100 µM. *ns*= no significant differences. Blue lines represent median values of fork velocity. (F) Schematic representation of incubation times with the chemical inhibitors and nucleoside analogues for the evaluation of replication speed and fork asymmetry. (G) Box and whiskers plot showing the replication fork asymmetry caused by mre11i or lack of TDP1 in NS and TDP1kd cells. Whiskers represent 10-90 percentiles. *ns*= no significant differences. *Lower panel:* examples of symmetric and asymmetric forks.

Based on our observation that TOP2α-mediated cleavage complexes could be stabilized by ETO ahead or behind of the ongoing replication fork (Fig. 3), we hypothesized that TOP2αcc would challenge the progression of the replication fork. To test the ability of MRE11 and TDP1 to remove these blocking proteins from DNA termini, we examined the replication fork progression in NS and TDP1kd cells by the DNA fiber assay. Cells pre-treated or not with mre11i were sequentially pulse-labeled for 15 min with IdU, followed by a 45-min pulse with CldU in the presence or absence of ETO (Fig. 4B). As shown in Fig. 4C-D, ETO significantly reduced the CldU/IdU ratio in both NS and TDP1kd cells (*P*<10^-4^). As expected, ETO-treated TDP1kd cells also showed significant differences compared to ETO-treated NS cells (P=0.006). However, mre11i pre-treated and ETO-treated TDP1kd cells showed a decreased replication forks progression that was similar to both the combined treatment in NS cells and the ETO-treated TDP1kd cells. Interestingly, both TDP1kd control and mre11i pre-treated NS cells showed a significant reduction (*P*<0.05) of the replication fork progression as compared to NS control cells.

Various DNA repair-deficient cells demonstrated to have reduced replication fork speed (Fu, Martin et al., 2015, Wilhelm, Ragu et al., 2016). The accumulation of endogenous DNA lesions can also lead to replication fork stalling, with sister forks starting from a single origin showing asymmetry. Thus, we first analyzed replication fork speed from NS and TDP1kd cells exposed or not to mre11i. As shown in Fig. 4E, the lack of TDP1 resulted in a diminished velocity of the replication fork (*P*=0.0002). However, the inhibition of MRE11 activity did not affect significantly the replication fork speed of NS cells. To analyze replication fork asymmetry, NS and TDP1kd cells were sequentially pulse-labeled with IdU and CldU for 20 min each, in the presence or not of mre11i (Fig. 4F). As shown in Fig. 4G, mre11i significantly increased replication fork asymmetry in NS cells compared to controls (P<0.05), while TDP1 deficiency alone did not promote sister forks asymmetries.

Overall, our findings suggest that TDP1 and MRE11 exhibit an epistatic interaction in removing ETO-induced TOP2αcc during S/G2 phases, facilitating the progression of the replication fork.

### TDP1 promotes DSB end-resection downstream to MRE11 activity

While classical non homologous end-joining typically requires unresected DSB to proceed, the other main DNA DSB repair pathways, microhomology-mediated end-joining and homologous recombination, require nucleolytic resection of 5’ ends to generate 3’ ssDNA overhangs (Patterson-Fortin & D’Andrea, 2020). To determine whether lack of TDP1 affect DSB end-resection, we analyzed ssDNA formation in S/G2 phase cells. With this aim, we examined ssDNA formation in CENP-F+ nuclei by both immunodetection of BrdU under native conditions and the ssDNA binding protein RPA.

Cell cultures were pre-treated or not with mre11i and treated with ETO by 1h and analyzed immediately or following 2h recovery in ETO-free media (Fig. 5A-E). As shown in Fig. 5A, a 1h treatment with ETO caused a slight increase in the BrdU signals in both NS and TDP1kd cells. However, BrdU signal intensity was higher after 2h of recovery. The mre11i pre-treatment of NS cells reduced significantly the intensity of BrdU induced by ETO to basal levels (*P*<10^-4^) at both times. Interestingly, ETO-treated TDP1kd cells after 2h of recovery showed an increased intensity of BrdU staining that was significantly lower (∼58%) than that found in NS cells (*P*<10^-4^). However, pre-treatment with mre11i of TDP1kd cells treated with ETO and 2h recovery diminished significantly the intensity of BrdU (*P*<10^-4^) returning them to basal levels. On the other hand, both NS and TDP1kd cells pre-treated with mre11i and treated with ETO did not show significant differences.

**Figure 5:**
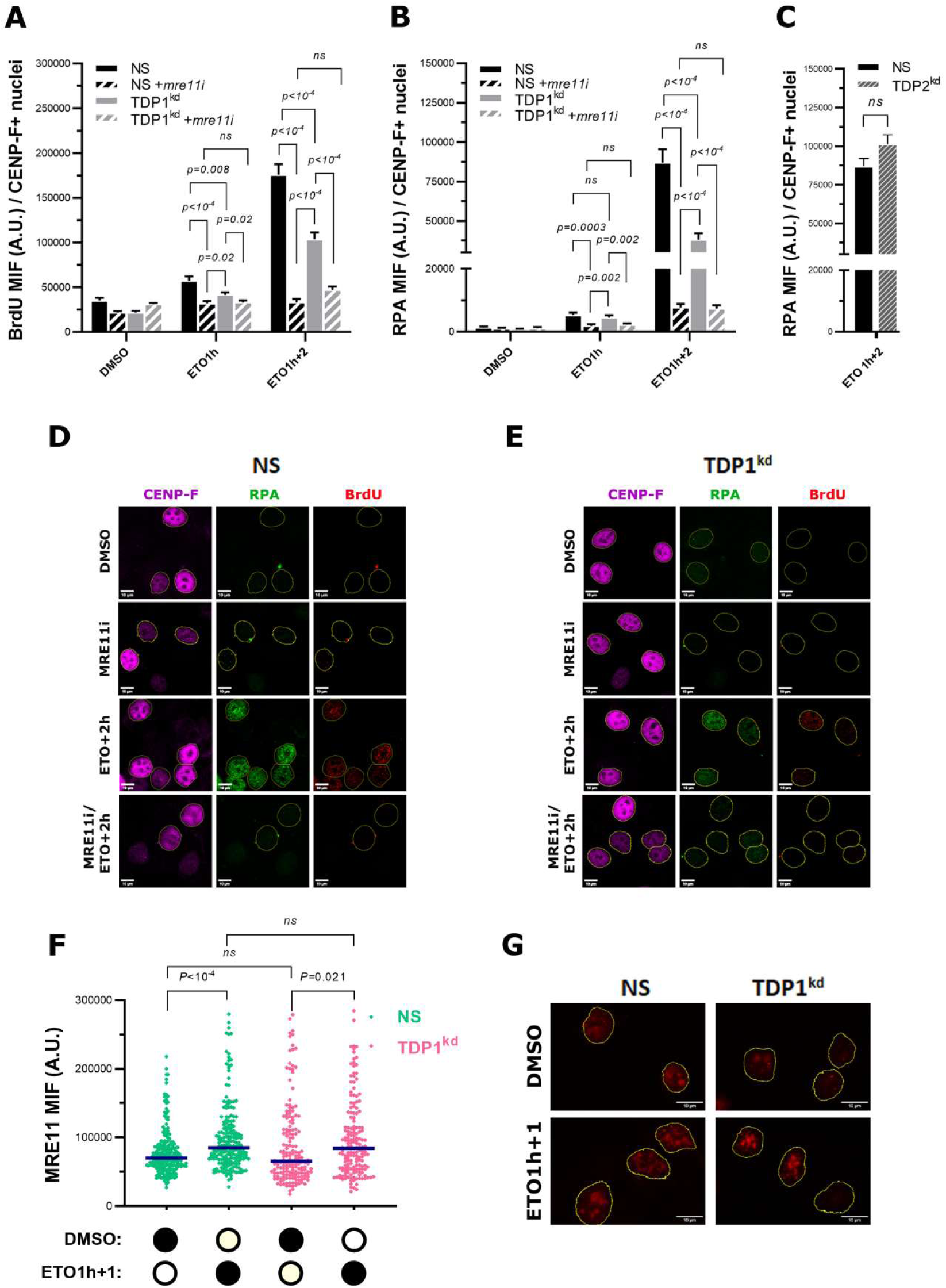
Lack of TDP1 impairs ssDNA formation during ETO-stimulated DSB end resection. NS, TDP1kd or TDP2kd cells were incubated with BrdU by 24 h, pre-treated or not with mre11i and then treated with ETO 17 µM by 1h and analyzed immediately or after 2 h of incubation in ETO-free culture media. The cells were then examined for BrdU and RPA immunolabeling under native conditions in the CENP-F+ population. (A) BrdU median intensity of fluorescence (MIF) in CENP-F+ nuclei of NS or TDP1kd cells. A.U.= arbitrary units. *ns*= no significant differences. (B) RPA median intensity of fluorescence (MIF) in CENP-F+ nuclei of NS and TDP1kd cells. *ns*= no significant differences. (C) RPA median intensity of fluorescence (MIF) in CENP-F+ nuclei of NS and TDP2kd cells. *ns*= no significant differences. (D) Maximal intensity projection images showing BrdU and RPA and CENP-F labeling of NS cells. (E) Maximal intensity projection images showing BrdU and RPA and CENP-F labeling of TDP1kd cells. (D-E) yellow lines show CENP-F+ nuclei. (F) MRE11 median intensity of fluorescence (MIF) was evaluated. Blue lines represent median values. (G) Representative images showing MRE11 chromatin loading.

The analysis of the ssDNA binding protein RPA showed a similar pattern (Fig. 5B, 5D and 5E). While 1h treatment with ETO caused a slight increase in RPA intensity, after 2h of recovery the intensity of RPA was much more evident in both NS and TDP1kd cells. Again, TDP1kd cells treated with ETO showed a lower intensity of RPA compared with NS cells (*P*<10^-4^). The intensity of RPA staining in TDP1kd cells represented ∼44% of the signals found in NS cells. Similarly, levels of RPA in both NS and TDP1kd cells under combined mre11i and ETO treatment were significantly lower (*P*<10^-4^) compared to ETO treatment alone. However, the combined treatment did not show significant differences between both cell lines. To determine whether the lower levels of ssDNA found were specific to TDP1kd cells; we analyzed the levels of RPA in TDP2kd cells. As depicted in Fig. 5C, the intensity of RPA staining in response to ETO was similar in both NS and TDP2kd cells.

To determine whether the reduced amount of ETO-induced ssDNA in the absence of TDP1 could be related to a limitation in the MRE11’s capacity of binding to chromatin, we analyzed the chromatin loading of MRE11 by immunofluorescence after protein extraction. As shown in Figure 5F-G, the treatment with ETO stimulated significantly the chromatin loading of MRE11 in both NS and TDP1kd cells (*P*<0.025). In addition, there were no significant differences between both cell lines, either in controls or after treatment with ETO.

Together, our results suggest that ssDNA tracks generated during DNA end-resection are promoted by TDP1 following stimulation with ETO in a MRE11-dependent manner.

### TDP1 promotes DNA nascent strand degradation during replicative stress

Stalled replication forks can undergo fork reversal, creating a “chicken foot” structure that contains a single end DSB that is vulnerable to nucleolytic degradation by a process known as nascent strand degradation (NSD). To determine whether lack of TDP1 impact on the NSD of DNA triggered by ETO at stalled forks, cells were incubated with BrdU by 30 min and treated with ETO 17 μM for 3h and subjected to DNA fiber analysis (Fig. 6A). As shown in Fig. 6B, ETO-treatment reduced BrdU track length in NS cells by 3.75-fold compared to controls (*P*<10^-4^). However, TDP1kd cells showed a milder effect (Fig. 6C), with a reduction of track lengths of 1.95-fold compared to controls (*P*<10^-4^). This corresponded to a 52% reduction in ETO-induced NSD in TDP1kd cells compared to NS cells. To determine if unstable reversed forks could also contribute to the reduced NSD capability in TDP1kd cells, we analyzed replication fork restart (suppl. Fig 2). The cells were labeled with IdU by 30 min, and then treated with ETO by 2 h followed by incubation with CldU by an additional 30 min (suppl. Fig 2A). As depicted in suppl. Fig. 2B, there were no differences in the restart of the replication forks between both NS and TDP1kd cells. However, ETO-treatment caused an increased number of stalled forks (*P*=0.015) and a reduced amount of new origin firings (*P*=0.013) in TDP1kd compared to NS cells.

**Figure 6:**
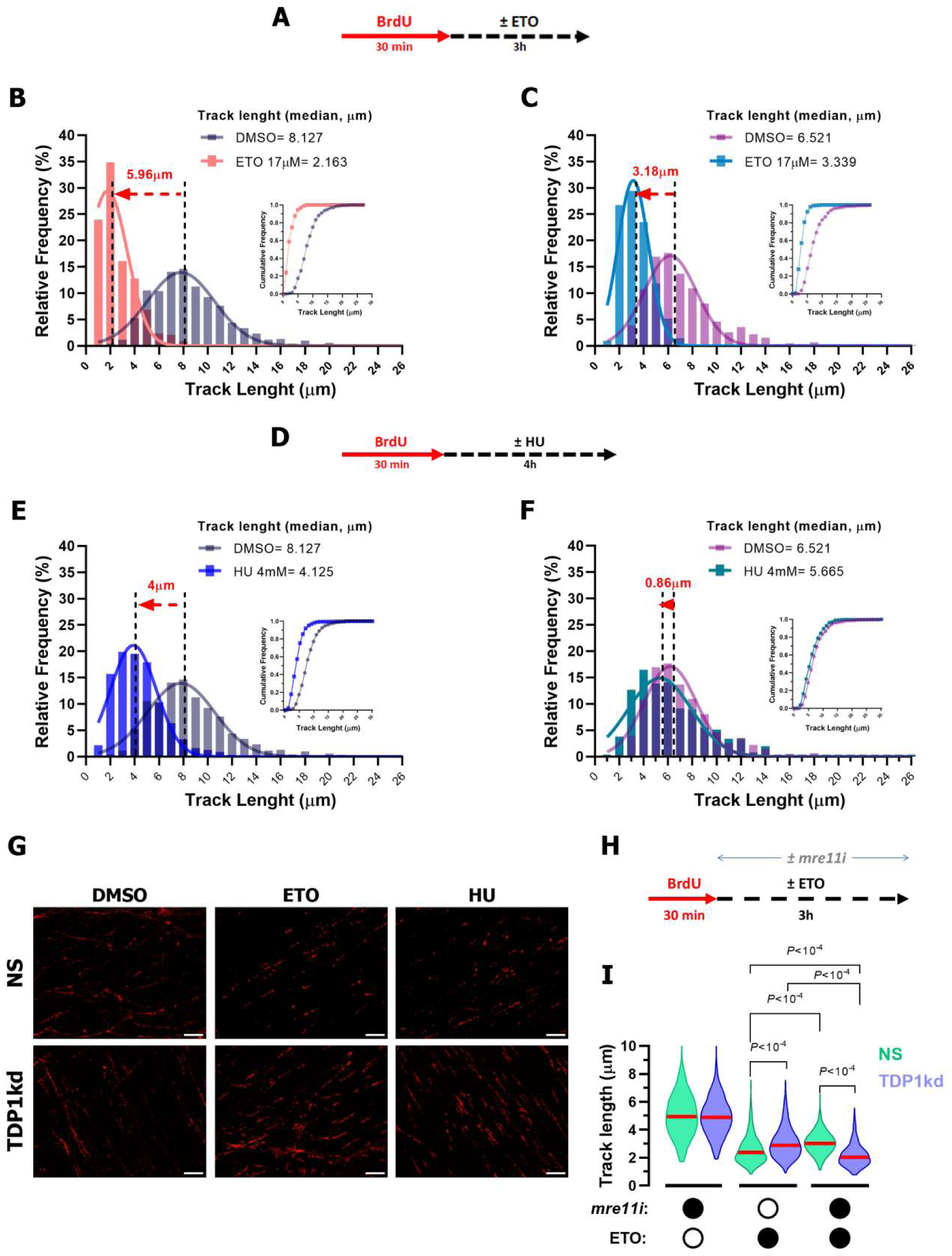
Loss of TDP1 diminishes nascent strand degradation at stalled replication forks. Degradation of newly synthesized DNA at stalled replication forks was evaluated at NS and TDP1kd cells following incubation replication stress-inducing agents. (A) Experimental timeline showing incubation with BrdU and ETO. (B) BrdU track length distribution of NS cells after exposure to ETO or DMSO. Inset shows cumulative frequency distribution. A total of 682 (control) or 422 (ETO) fibers were scored from 4 independent experiments. (C) BrdU track length distribution of TDP1kd cells after exposure to ETO or DMSO. Inset shows cumulative frequency distribution. A total of 584 (control) or 404 (ETO) fibers were scored from 4 independent experiments. (D) Experimental timeline showing incubation with BrdU and HU. (E) BrdU track length distribution of NS cells after exposure to HU or DMSO. Inset shows cumulative frequency distribution. A total of 682 (control) or 547 (HU) fibers were scored from 4 independent experiments. (F) BrdU track length distribution of TDP1kd cells after exposure to HU or DMSO. Inset show cumulative frequency distribution. A total of 584 (control) or 504 (HU) fibers were scored from 4 independent experiments. (G) Representative DNA fiber images. Scale bar, 10 µm. (H) Experimental timeline showing incubation with BrdU and ETO with or without mre11i. (I) Violin plot showing BrdU track length of DNA fibers from ETO-exposed NS and TDP1kd cells pretreated or not with mre11i. Red lines represent median values. From 500 to 800 fibers were scored and data came from 5 independent experiments.

Then, we analyzed whether the reduced NSD observed in the absence of TDP1 is also evident following incubation with other agent known to induce nascent strand degradation of reverted forks such as the ribonucleotide reductase inhibitor hydroxyurea (HU) (Thangavel, Berti et al., 2015). We cultured cells in the presence of BrdU by 30 min and then treated with a high-dose of HU (4mM) by 4h before the DNA fiber analysis (Fig. 6D). As shown in Fig. 6E, treatment with HU in NS cells caused fiber lengths 1.97-fold shorter as compared to control cultures (*P*<10^-4^). Nevertheless, HU treatment in TDP1kd cells (Fig. 6F-G) revealed a less pronounced decrease in track lengths that was 1.15-fold lower as compared to control cultures (*P*<10^-4^). Thus, the NSD triggered by HU in TDP1kd cells signified a reduction of 58% compared to that obtained in NS cells.

To determine if there is a functional interaction between TDP1 and MRE11 on NSD, we performed a DNA fiber analysis in the presence of a MRE11 inhibitor (MRE11i) and ETO (Fig. 6H-I). As shown in Fig. 6I, inhibiting the MRE11 exonuclease activity during ETO exposure increased fiber track length compared to ETO alone (*P*<10^-4^). The lack of TDP1 also caused increased fiber track lengths as compared to ETO in NS cells (*P*<10^-4^). However, fiber tracks were significantly shorter in MRE11i+ETO-exposed TDP1kd cells compared to ETO alone, MRE11i+ETO treated NS, and ETO-exposed NS cell groups (*P*<10^−4^).

These data strongly suggest that the absence of TDP1 alters the extent of NSD independently of MRE11 activity, regardless of the cause of replication fork stalling.

### Lack of TDP1 combined with the inhibition of MRE11 do not increase the sensitivity to ETO

In order to determine whether the sensitivity to ETO is affected by the combined inhibition of TDP1 and MRE11, we analyzed the cell viability of NS and TDP1kd cells in the presence or not of mre11i (20 μM). The curves in Fig. 7A represent the viability of the different cell lines after 96 h of exposure to different concentrations of ETO. In Fig. 7B is represented a table with the IC50 estimated for each condition. As expected, the pre-treatment with mre11i in NS cells showed an IC50 significantly lower (P<0.05) than NS cells. Similarly, TDP1kd cells showed hypersensitivity to ETO (*P*<0.05) compared to NS cells. The pre-treatment with mre11i of TDP1kd cells also showed an IC50 that was lower than NS cells (*P*<0.05). However, it was not significantly different from both NS pre-treated with mre11i or TDP1kd cells (Fig. 7B).

**Figure 7:**
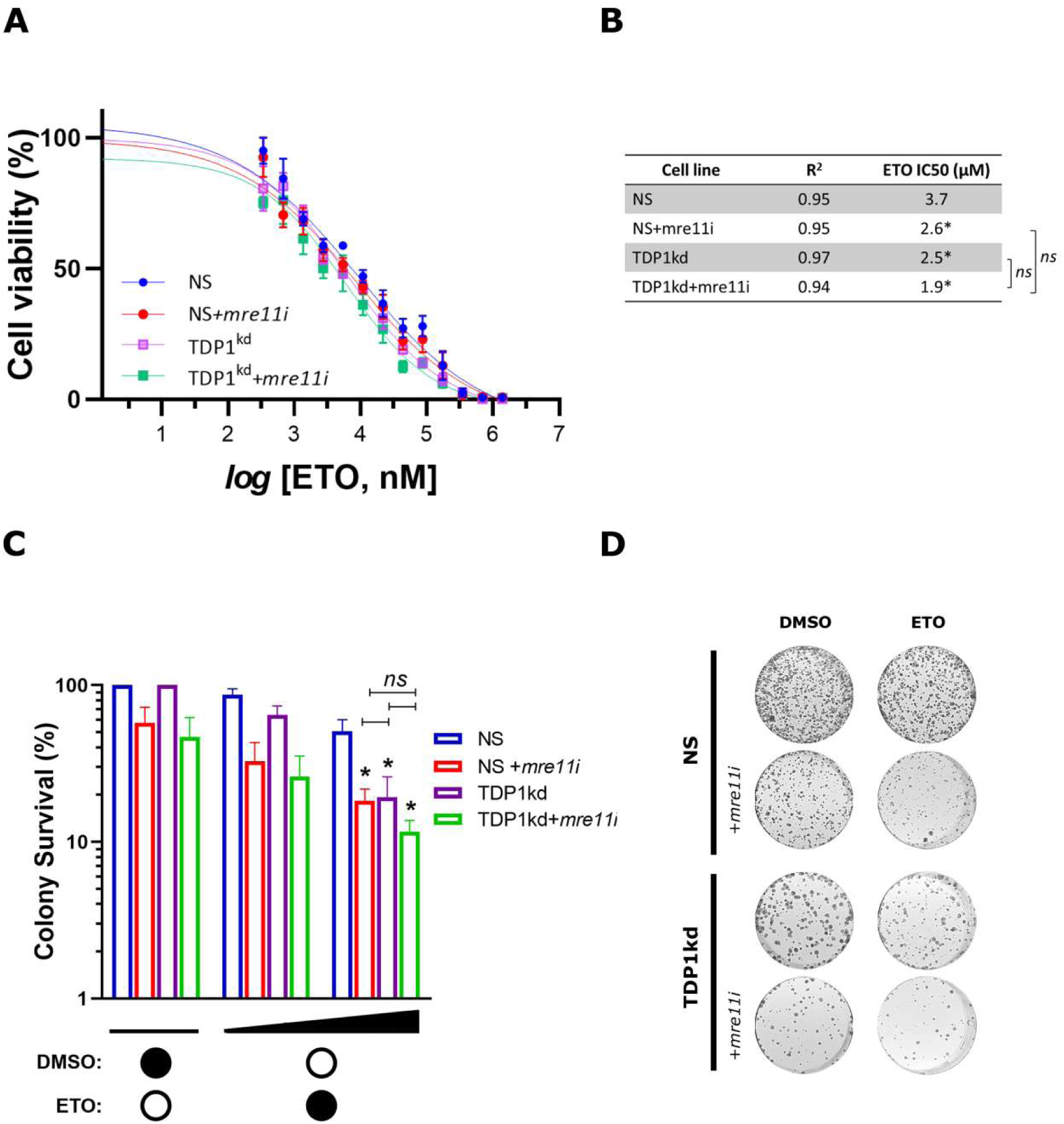
TDP1 and MRE11 act epistatically to promote survival in ETO-treated cells. (A) Cell viability was assessed in NS and TDP1kd cells pretreated with or without 20 µM mre11i, followed by 20 h of ETO treatment, and measured 96 h after ETO exposure. Error bars represent s.e.m. (B) Table of ETO IC50 values. \**P* < 0.05; *ns*, not significant differences. (C) Colony formation assay. Cells were pretreated with 50 µM mre11i and then exposed to 0.05 or 0.2 µg/ml ETO for 20 h. Colonies were counted after 12–14 days. Error bars represent s.e.m. \**P* < 0.05; ns, no significant differences. (D) Representative images of the colony formation assay. Treatment with 0.2 µg/ml ETO is exemplified.

To further corroborate these results, we performed a colony formation assay (Fig. 7C and 7D) using a higher dose of mre11i (50 μM). NS or TDP1kd cells, pre-treated or not with mre11i, were treated or not with sub-lethal doses of ETO and analyzed after 12-14 days. As seen in Fig. 7C-D, NS cells pre-treated with mre11i and TDP1kd cells pre-treated or not with mre11i were hypersensitive to the higher dose of ETO compared to their respective controls (*P*<0.05). However, TDP1kd cells pre-treated with mre11i and treated with ETO didńt show significant differences with TDP1kd or NS cells pre-treated with mre11i and treated with ETO.

Overall, our findings support an epistatic relationship between TDP1 and MRE11 in response to ETO treatment.

## DISCUSION

Defective removal of TOP2cc can result in severe genomic and cellular consequences, including the accumulation of persistent DSB that leads to mutagenesis or cell death. In this study, we aimed to explore the involvement of TDP1 in the removal of drug-stabilized TOP2αcc in human cells and its functional relationship with the nuclease MRE11. TOP2αcc stabilized by ETO accumulated throughout the cell cycle, being differentially increased in the S/G2 phases of TDP1kd cells compared to NS cells. In addition, there was no reduction of γH2AX levels in either the S or G2 phases in cells lacking TDP1. The increased accumulation of ETO stabilized-TOP2αcc demonstrates an impaired ability of TDP1-deficient cells to resolve them during the replicative and post-replicative phases of the cell cycle. The involvement of TDP1 in replication-associated DNA repair has been previously suggested in studies addressing camptothecin-induced TOP1-mediated cleavage complexes (Ryan, Squires et al., 1994), as well as during the removal of chain-terminator nucleoside analogs (Huang, Murai et al., 2013).

Topological stress resulting from the advance of RNA polymerase II requires DNA topoisomerases to reverse accumulated DNA supercoiling to allow transcription elongation. The binding sites of TOP2A and TOP2B to DNA are enriched at regulatory sequences and gene bodies of transcriptionally active genes. These sites correlate with RNA polymerase II occupancy and DNAse I hypersensitivity (Canela, Maman et al., 2019, Dellino, Palluzzi et al., 2019, Martinez-Garcia, Garcia-Torres et al., 2021, Thakurela, Garding et al., 2013, Yu, Davenport et al., 2017). In addition, the distribution of TOP2-associated DNA DSBs was reported to be augmented at promoters; 5’ splice sites, and active enhancers (Dellino et al., 2019, Singh, Szlachta et al., 2020). Thus, we analyzed the contribution of the transcription machinery collisions with TOP2αcc to generate DNA DSB. Curiously, the inhibition of transcription elongation did not raise substantially the accumulated TOP2αcc levels, as would be expected by reducing the likelihood of collisions. Moreover, the γH2AX signals that would result from the collision of the transcription machinery with either TOP2α or TOP2β-mediated cleavage complexes did not diminish by the presence of the transcription inhibitors. Our results demonstrate that, in our experimental conditions, ETO-induced DNA DSBs are not attributable to transcription machinery collisions with TOP2-mediated cleavage complexes, regardless of TDP1 presence. Similar results were reported previously (Gomez-Herreros, Zagnoli-Vieira et al., 2017), even after incubating with ETO for twice the time analyzed here, which corresponds to four times the global half-life of the ETO-induced TOP2αcc. In addition, they are in line with studies where exposure to TOP2 poisons did not alter the transcription levels of immediate early genes (Herrero-Ruiz, Martinez-Garcia et al., 2021). The accumulation of ETO-induced γH2AX showed an inverse correlation with TOP2αcc levels. In this regard, the highest levels of γH2AX were observed in the G1 phase, where the half-life of ETO-induced TOP2αcc was shortest. These results further demonstrate that the processing of TOP2αcc occurs with different kinetics throughout the cell cycle, suggesting different regulatory mechanisms.

TOP2 poisons cytotoxic effects involve targeting TOP2α during DNA replication (Burgess, Doles et al., 2008, Fan et al., 2008, Tammaro et al., 2013). As ETO-stabilized TOP2αcc would be responsible for the inhibition of the replication, we aimed to determine whether they form ahead or behind of the replication forks. We demonstrated in human cells that ETO-induced TOP2αcc forms on both sides of replication forks, suggesting a dual mechanism of replication inhibition involving both fork collapse and stalling. Previous studies analyzing plasmid DNA replication intermediates in *Xenopus* egg extracts, under TOP2 poisoning conditions, revealed that TOP2α acts behind replication forks (Heintzman, Campos et al., 2019, Lucas, Germe et al., 2001, Van Ravenstein, Mehta et al., 2022). One of them raised the prospect that ETO might occasionally trap TOP2α ahead of the forks (Lucas et al., 2001). In this sense, various conditions may contribute to the accumulation of positive supercoiling ahead of replication forks, including regulation problems of replication fork rotation (Schalbetter, Mansoubi et al., 2015), transcription-stimulated and cohesin-mediated chromatin loop extrusion (Benedetti, Racko et al., 2017), head on replication-transcription conflicts (Browning & Merrikh, 2024), and certain external stimuli, such as heat-induced stress (Roti Roti & Painter, 1982). A recent study using new probes for assessing the topological state of the DNA, has also been able to detect the accumulation of positive supercoiling at telomeres and fragile sites after both replicative stress and chemical inhibition of TOP2 (Ghilain, Vidal-Cruchez et al., 2024).

MRE11 is an essential protein for the removal of TOP2cc, as its absence or mutations affecting its endonuclease activity lead to the accumulation of spontaneous TOP2cc (Hoa, Shimizu et al., 2016a). Here we found an epistatic relationship between MRE11 and TDP1 in the removal of ETO-induced TOP2αcc during S/G2 phases of the cell cycle. Further support came from the analysis of the replication fork stalling after exposure to ETO, which revealed defects consistent with this functional interaction. Notably, both the lack of TDP1 and the inhibition of the MRE11’s nuclease activity disrupted the normal replication fork dynamics. It is noteworthy that the mre11i used in this study was reported to inhibit the exonuclease activity of MRE11 (Dupre, Boyer-Chatenet et al., 2008) but also to partially inhibit its endonuclease activity in vitro (Deshpande et al., 2016).

Unrestricted DSB end resection and stalled replication forks lead to genome instability. MRE11 is the main nuclease in the step of short-range DSB end resection. Thus, we investigated whether TDP1 conditions the MRE11-mediated DSB end resection process. We found that DNA ends resection was constrained by lack of TDP1 during the S/G2 phases of the cell cycle. As TDP1 directly removes single protein adducts or nucleotides from DNA ends (Huang et al., 2013), it might also promote end resection by an indirect way. We demonstrated here that the exonuclease activity stimulated by TDP1 acts downstream of the MRE11 activity.

ETO was reported to induce replication fork reversal and DNA NSD (Zellweger, Dalcher et al., 2015). Therefore, we next explored the involvement of TDP1 in NSD. We found that TDP1 deficiency impaired the NSD response to the replication fork stalling triggered by either TOP2-blocked DNA ends or nucleotide pool depletion. In addition, we observed that the NSD promoted by TDP1 was independent of the nuclease activity of MRE11. Interestingly, the inhibition of MRE11’s nuclease activity in a TDP1-deficient context stimulated the NSD induced by ETO, probably through the activity of other nucleases. In this regard, late S-phase replication forks have an impaired protection of stalled forks and are susceptible to NSD by an EXO1-dependent process (Colicino-Murbach, Hathaway et al., 2024). Our findings also reinforce the idea that DNA end processing is controlled by distinct regulatory mechanisms during DSB end resection and at stalled replication forks, as suggested previously (Zhao, Hou et al., 2023).

To validate the epistatic relationship between TDP1 and MRE11 in response to ETO, we assessed the viability of cells deficient in either or both protein activities. Under both short– and long-term culture conditions, the single and double deficient cells displayed similar hypersensitivity to ETO, supporting an epistatic effect. In a *S. cerevisiae* strain lacking Tdp1 (tdp1Δ), the expression of a nuclease-dead Mre11 mutant was reported to increase the sensitivity to ETO (Hamilton & Maizels, 2010). This suggests that, in yeast, Mre11 and Tdp1 function in parallel pathways against TOP2 poisons. Similarly, a previous study reported a synergism between MRE11 and TDP2 in response to ETO in human TK6 cells (Hoa et al., 2016a), indicating that both activities operate by different mechanisms against ETO-induced TOP2cc.

Overall, we demonstrate that TDP1 facilitates the replication-coupled removal of TOP2 poisons-stabilized TOP2αcc by promoting DNA end processing via both MRE11-dependent and –independent pathways. These findings support previous reports describing redundant roles for TDP1 and MRE11 nuclease activity in the replication-dependent removal of chain-terminating nucleoside analogs in vertebrate cells (Mohiuddin, Rahman et al., 2019). Collectively, our work reveals that beyond its known role in directly hydrolyzing TOP2αcc, TDP1 plays an indirect role in stimulating the nucleolytic excision of these protein adducts. It will be interesting to further dissect the complete scenario of TDP1’s interacting proteins through which it can promote the activity of DNA end processing after replicative stresses.

## METHODS

### Cell culture and chemical inhibitors

The HeLa cell lines were grown in RPMI 1640 (#31800022, Gibco) supplemented with 10% FBS (#FRA500, Internegocios) and maintained at 37°C with 5% CO_2_ in a humidified atmosphere. HeLa non-silencing control (NS), TDP1 (TDP1kd) and RAD21 (Rad21kd) knocked down cell lines were previously described (Borda et al., 2015, Kramar J., 2023). HeLa TDP2kd cells were generated by stable transfection using a pre-designed pRNAi-H1-hyg linearized plasmid (#SORT-A04, Biosettia) and Lipofectamine 2000 (#11668019, Invitrogen) according manufacturer’s instructions. After 48 h, selection was carried out in selection media containing 200 μg/ml hygromycin (#ant-hg-1, InvivoGen), followed by a 4-week clonal selection process. The TDP1kd and TDP2kd selected clones were verified by qRT-PCR to reduce the expression of their targets more than 80%. Rad21kd cell line was reduced in 68% (suppl. Figure 1). All the cell cultures were routinely monitored for mycoplasma contamination. The following inhibitors were used where indicated: Etoposide (ETO) (#E1383, Sigma-Aldrich), 5,6-dichloro-1-beta-D-ribofuranosylbenzimidazole (DRB) (#D1916, Sigma-Aldrich) and Mirin (mre11i) (#sc-203144, Santa Cruz), Aphidicolin (APH)(# 178273, Sigma-Aldrich), Hydroxyurea (HU, Sigma-Aldrich).

### TOP2αcc and γH2AX analysis by flow cytometry

The evaluation of drug-induced TOP2αcc by flow cytometry was performed as reported previously (de Campos-Nebel, Palmitelli et al., 2016, de Campos Nebel, Palmitelli et al., 2017). Briefly, harvested cell cultures treated with either DMSO or ETO (with or without Mirin pre-treatment), were extracted in PHEM buffer (65 mM PIPES, 30 mM HEPES, 10 mM EGTA, 2 mM Mg_2_Cl, pH 6.9) containing 0.5% Triton-X100, Heparin (100U) and 1 mM PMSF under rotatory mixer by 5 min. The samples were fixed in 1% paraformaldehyde by 30 min. Following a 1 h blocking step (3% BSA and 0.5% Triton X-100 in PBS), samples were incubated with a primary antibody anti-TOP2a antibody (#sc-3659, Santa Cruz) in blocking buffer for 2h. After washing with phosphate buffered saline (PBS), samples were incubated with Alexa Fluor 488-conjugated goat anti-mouse antibody (#A11001, Thermo Fisher) in blocking buffer by 1h. Samples were then washed with PBS and stained with 200 μg/ml RNAse A and 20 μg/ml propidium iodide in PBS.

To evaluate the γH2AX formation, cell cultures were harvested and washed with PBS. They were fixed with 90% of methanol by 30 min at –20°C and washed in PBS. They were then permeabilized in 0.25% Triton X-100 solution in PBS by 5 min at room temperature. Samples were blocked and incubated with a primary antibody anti-γH2AX (#05-636-I, Millipore), followed by incubation with Alexa Fluor 488-conjugated goat anti-mouse antibody (#A11001, Thermo Fisher) and incubation with RNAse A and propidium iodide, as mentioned above.

In both cases, at least 25,000 cells/treatment were analyzed from 3 to 6 independent experiments.

### Single-molecule analysis of DNA replication

To analyze the replication fork progression, cells were first labeled with 20 μM IdU (#sc-205720, Santa Cruz) for 15 min, followed by 100 μM CldU (#C6891, Sigma-Aldrich) for a further 45 min in the presence of DMSO or ETO. For cultures pre-treated with 100 μM mre11i, the drug was present throughout the incubations with both halogenated analogs. To analyze fork assymetry, the cells were incubated with IdU and CldU as above but for 20 min each. The cells were harvested by trypsinization and then, labeled cells were loaded onto a slide. A cell lysis solution (200 mM Tris–HCl, pH 7.5, 50 mM EDTA and 0.5% SDS) was added to lyse the cells for 10 min. The slides were angled to allow the spread of the DNA for 5 min. After air drying, the fibers were fixed by 10 min in a 3:1 (vol/vol) methanol: acetic acid solution and denatured in 2.5 M HCl at room temperature by 60 min. Then, the slides were washed twice with 1 X PBS and blocked with blocking solution (5% goat serum, 0.1% Triton X-100) before adding anti-BrdU antibodies (#ab6326, Abcam and #347580, BD) and corresponding FITC– and Cy3-labeled secondary antibodies (#PA1-29948 and #A10521, Thermo Fisher). Images of the slides were captured under an Olympus FV1000 confocal microscope and analyzed from 3 to 4 independent experiments using the FiberQ software (Ghesquiere, Elsherbiny et al., 2019).

For the analysis of nascent strand degradation (NSD), cells were labeled with 32 μM of BrdU (#B5002, Sigma-Aldrich) for 30 min, washed and then treated with DMSO, ETO or HU as indicated. Cells were harvested and processed as described previously. Images were captured under the confocal microscope and the fibers length was measured from 3 to 4 independent experiments with the ImageJ software.

### Single-stranded DNA (ssDNA) formation and MRE11 chromatin loading by immunofluorescence staining

The analysis of ssDNA formation in S/G2 phases was carried out as described previously (Mukherjee, Tomimatsu et al., 2015) with minor modifications. Briefly, cells were grown in the presence of 10 μM BrdU (#B5002, Sigma-Aldrich) for 24 h, then pre-treated or not with mre11i and treated with DMSO or ETO as indicated. The cells were rinsed with cold PBS, incubated by 10 min with pre-extraction buffer (100 mM PIPES pH 7.5, 100 mM NaCl, 1 mM EDTA, 3 mM MgCl_2_, 300 mM Sucrose, 0.5 % Triton X-100) at 4°C, and incubated 5 min in CSK buffer (10 mM Tris-HCl pH 7.4, 10 mM NaCl, 3 mM MgCl_2_, 1mM EDTA, 0.5% Triton X-100, 0.5% Na-Deoxycholate) at 4°C prior to fixation by 20 min in 4 % paraformaldehyde. Permeabilization was done by 10 min with 0.5 % Triton X-100. The samples were then incubated in blocking solution containing 5 % bovine fetal serum in PBS by 1h, and then with anti-BrdU (#sc-32323, Santa Cruz), anti-RPA (#2208S, Cell Signaling), and anti-CENP-F (#sc-22791, Santa Cruz) antibodies. Samples were incubated with the respective fluorophore-conjugated secondary antibodies (#A11020, Thermo Fisher; #sc-2991, Santa Cruz; #A21245, Thermo Fisher) by 1h.

For MRE11 immunofluorescence staining, cells were pre-extracted with 0.5% Triton X-100 for 5 min to release the soluble fraction of MRE11, followed by fixation with 4% paraformaldehyde for 20 min. After blocking with 5% BFS for 1 h, anti-MRE11 antibodies (#4847, Cell Signaling) were incubated for 2 h at room temperature, followed by 1 h incubation with secondary antibodies (#A11072, Thermo Fisher). Finally, cells were mounted with Vectashield antifade mounting media with DAPI (#H-1200, Vector Laboratories).

Images from 3 independent experiments were acquired using an Olympus FV1000 confocal microscope and analyzed with ImageJ software.

### Cell viability

Cell viability was determined by the standard MTT (3-(4,5-dimethylthiazol-2-yl)-2,5-diphenyltetrazolium bromide) (#475989, Sigma-Aldrich) reduction assay. NS or TDP1kd cells (2.5×10^3^ per well) were seeded into 96-well plates and incubated overnight at 37°C. Different doses of ETO were added for 20 h in the presence or not of Mirin (20 μM). After drug treatment, cells were washed twice in PBS and incubated in fresh medium. After 96 h, MTT solution (1 mg/ml) was added to each well and incubated by 3 h in the dark at 37°C. The supernatant was removed and, after washing, 100 μl DMSO was added. The absorbance of each sample was measured at 550 nm using a microplate reader. Cell survival was calculated by normalizing the absorbance of the treated samples to the absorbance of the DMSO-treated controls. Survival curves were generated using a non-linear regression model, and the relative IC50 was calculated using the Graphpad Prism 9.5 software. Four independent experiments were performed in triplicates.

### Clonogenic survival

HeLa NS cells (7.5×10^2^) or TDP1kd cells (5.0×10^2^) were seeded on 6-well tissue culture dishes. After 20 h, the cells were pre-treated or not with mre11i (Mirin, 50 μM), and treated with different doses of ETO by 24 h. After washing, the colonies were grown for 12-14 days. Clusters containing at least 50 cells were considered colonies. The colonies were washed twice with 1X PBS, fixed with 100% methanol, and stained with 0.5% crystal violet solution. Colony survival upon drug treatment was determined by normalizing the total number of colonies formed with plating efficiency under individual conditions. Data were obtained from 3 to 4 independent experiments. Graphpad Prism 9.5 software was used to graph the results.

### Statistical Analysis

Statistical analysis was performed with Graphpad Prism software. Data from TOP2αcc were analyzed by one-way ANOVA or paired comparisons by *t*-tests. γH2AX immunofluorescence and MRE11 chromatin loading were analyzed with a Kruskal-Wallis test. The replication fork progression, the frequency distribution of DNA tracks length and the ssDNA formation were analyzed with the Mann-Whitney *U* test or Kruskal-Wallis test for multiple comparisons.

## ACKNOWLEDGEMENTS

We thank Federico Fuentes (IMEX) for technical support with microscopy. This work was supported by grants from the National Scientific and Technical Research Council (CONICET, PIP-21-0886] from Argentine to MdCN.

## AUTHOR CONTRIBUTIONS

**Néstor Aznar:** Conceptualization; Formal analysis; Investigation; Methodology; Visualization; Validation; Writing—review and editing. **María Camila Gosso:** Conceptualization; Investigation; Methodology; Writing—review and editing. **Irene Larripa:** Supervision; Writing—review and editing. **Marcela González-Cid:** Conceptualization; Supervision; Writing—review and editing. **Marcelo de Campos Nebel:** Conceptualization; Formal analysis; Investigation; Validation; Supervision; Funding acquisition; Writing—original draft.

